# The near-normal viability of heterozygote for a lethal enhancer allele suggests a route for *cis*-regulatory evolution

**DOI:** 10.1101/2023.09.30.560335

**Authors:** Michael Z. Ludwig

## Abstract

A variety of evolutionary processes shape the structure of *cis*-regulatory elements such as enhancers. Functionally important regulatory sequences tend to be conserved as result of negative selection against deleterious mutations and positive selection for better-canalized performance. However, some forms of stabilizing selection can maintain functional conservation of *cis*-regulatory elements for long periods of evolutionary time despite structural transcription binding sites turnover (Ludwig et al, 1998). In addition, compensatory evolution can even accelerate the substitution process in large populations to level greater than the neutral rate of substitution (Carter and Wagner, 2002). The effects of lethal mutations in *cis*-regulatory regions on viability in heterozygotes and the impact of such mutations on evolutionary processes have not been substantially addressed.

We ask a fundamental biological question: How does a lethal mutant in an enhancer region affect viability when carried over a dominant “normal” allele?

Here *cis*-regulatory mutation ***eve***^***ΔMSE***^ of *even-skipped* gene (***eve)*** in *D. melanogaster*, that is lethal in homozygote, has been tested for relative viability in heterozygous state. In ***eve***^***ΔMSE***^ the 480-bp fragment corresponding to MSE (Minimal Stipe Element) of the ***eve*** stripe 2 enhancer was deleted and replaced with an unrelated DNA fragment (containing *white*^*+*^ gene marker) by ends-out homologous recombination (Ludwig et al, 2011). We discovered that relative viability of flies carrying *cis*-regulatory mutation ***eve***^***ΔMSE***^ in heterozygote was not reduced compared to the viability of wild-type flies. In contrast, the viability of heterozygotes carrying a homozygote-lethal nonsense ***eve***^***R13***^ mutation (Fujioka et al, 1999) was significantly impacted. Our explanation to this surprising phenomenon relays on action of ***eve*** transcriptional autoregulation. The transcription factor (TF) EVE regulates its own expression level through the autoregulatory *cis*-regulatory module. It could elevate the expression level from one dose to the level sufficient to restore up to 100% fitness in the heterozygote with ***eve***^***ΔMSE***^ lethal allele.

This example suggests that some *cis*-regulatory mutants (including the deleterious ones) may exist in populations as heterozygotes with high fitness for many generations, before possibly encountering an epistatic interaction with a compensatory mutation at a different site in the genome. Our study is consistent with the compensatory mechanisms of evolution for functionally important *cis*-regulatory elements.

## Introduction

Many functionally important noncoding sequences in genomes are not evolutionary conserved (Ludwig and Kreitman, 1995; Ludwig, 2002). Some of these have lineage-specific functions and others are simply missing by comparative genomics methods. There are many examples of extensively diverged regulatory sequences that have retained expression specificity (Ludwig et al, 1998). What the evolutionary processes governing the changes in these functionally conserved elements?

From experimental studies of evolutionary changes in functionally conserved regulatory elements in different species, the following observations can be gleaned: there is a lack of complete conservation of functional TF binding sites; there is rapid turnover of spacer sequences between binding sites; conservation is apparent; turnover of binding site architecture occurs; and there is co-evolution of TF binding sites within an enhancer.

The stripe 2 enhancer (S2E) from segmentation gene ***eve*** in *Drosophila* illustrates these points (Ludwig et al, 1998; Ludwig et al, 2000). This *cis*-element drives the expression of ***eve*** gene in a transverse stripe approximately 3 to 5 cells in wide in early blastula embryo of many different *Drosophila* species (Stanojevic et al, 1991; Small et al, 1992; Ludwig et al, 1998; Ludwig et al, 2005). S2E enhancer has been extremely well characterized in the fruit fly, *D. melanogaster*. For this reason, this *cis-*regulatory element has been intensively studied from an evolutionary perspective among other species, including Sepsid fly, *Themira putris* (Hare et al,2008). Enhancers from these two species, *melanogaster* and *putris*, which diverged about 100 million years ago, are unalignable at the DNA sequence level. Predicted locations of the functionally important transcription binding sites are different. However, in transgenic experiments a Sepsid *eve* S2E sequence was identified, as an element that drives the expression of a reporter gene in an embryonic transverse stripe in nearly the same location as observed with the endogenous *melanogaster* S2E enhancer (Hare et al, 2008).

A number of evolutionary models have been proposed to explain the observations concerning evolutionary changes in *cis*-elements such as enhancers. The presence of multiple TF binding sites naturally leads to the idea that the ‘model of stabilizing selection’ can be considered as a major mode of enhancer evolution (Ludwig, 2002). Applied to enhancer evolution, stabilizing selection can accommodate binding site turnover without disruption of primary enhancer function. The results of studies of the ***eve*** S2E evolution from *Drosophila* are consistent with the stabilizing selection model (Ludwig et al, 1998; Ludwig et al, 2000).

The evolution of some regulatory elements might also be consistent with a ‘model of compensatory selection’. A pair of mutations at different sites (loci) that are singly deleterious but restore normal fitness in combination may be called compensatory neutral mutations. Kimura (1985) demonstrated that these mutations could easily become fixed in the population by genetic drift when the genes are tightly linked. Carter and Wagner (2002) applied this model to the evolution of regulatory sequences. They found that large population size accelerates compensatory evolution, whereas small population sizes inhibit this form of drift from occurring.

However, the theoretical models listed above are not able to explain all aspects of *Drosophila* ***eve*** gene regulatory evolution. The following facts could complicate and challenge the feasibility of proposed models for evolution in S2E enhancer.

1. Both models operate for deleterious or lethal mutations. However, the transcription factors such as ***eve*** could be haploinsufficient genes. So, it is necessary to demonstrate that heterozygote with a lethal mutation for the TF at least could be alive.
2. The regulation of ***eve*** expression is extremely complex. The ***eve*** locus contains many *cis-*regulatory elements as enhancers, silencers, insulators, including some autoregulatory enhancers in the ***eve*** locus (Jiang et al, 1991; Small et al, 1996; Sarkison et al, 2000; Fujioka et al, 1999). The architecture of regulatory network is adjusted by natural selection to ensure robust gene expression. How does a mutation in one regulatory element will affect the functioning of the whole regulatory network?
3. The ***eve*** is dosage sensitive gene. In a transgenic rescue assay, one copy of a transgenic whole ***eve*** locus does not rescue the lethality of ***eve*** ^***R13***^ /***eve*** ^***R13***^ mutant with same success as two copies do (Ludwig et al, 2011).
4. In a population, any deleterious mutation has a better chance to be compensated by other *cis*or *trans*-mutation if it exists in substantially large numbers during many generations. According to theory, recessive lethal alleles could exist in large numbers in a population because of genetic drift, linkage with favorable alleles, and/or heterozygote advantage.

Herein, we addressed several questions pertaining to the regulatory evolution of the ***eve***.

First, we asked how does a lethal mutation in the ***eve*** S2E enhancer affect the organism’s viability when carried over a “normal” allele in heterozygote of *D. melanogaster*? Second, we investigated how does lethal mutation in S2E enhancer affect segmentation embryonic pattern in the heterozygote?

Third, we compare the viability of heterozygotes of lethal ***eve*** ^***R13***^ and ***eve*** ^***ΔMSE***^ mutants. The ***eve*** ^***R13***^ is a coding region lethal mutation that truncates the EVE protein within the homeodomain (Fujioka et al, 1999).

## Results

We herein study two ***eve*** lethal mutants. The ***eve*** ^***R13***^ is a coding region nonsense lethal mutation that truncates the protein within the homeodomain (Fujioka et al, 1999). The ***eve*** ^***ΔMSE***^ is *cis*-regulatory lethal mutant, where the 480-bp fragment corresponding to MSE (Minimal Stripe Element; see Small et al, 1992) of the ***eve*** stripe 2 enhancer was deleted and was replaced with the ***white***^***+***^ gene by ends-out homologous recombination (Ludwig et al, 2011).

### Relative Viability

We measure and compare two ***eve*** lethal mutants (***eve*** ^***R13***^ and ***eve*** ^***ΔMSE***^) relative viability in heterozygous state. The effects on both females and males heterozygous for lethal was compared to females and males not carrying a lethal (viability of these = 1). We demonstrated that viability of the ***eve*** ^***ΔMSE***^ lethal heterozygote is equal to that of the normal homozygote. The relative viability of female and male heterozygotes for ***eve*** ^***ΔMSE***^ were 1.0 and 1.04 correspondingly (Figure1A). Viability values were not significantly different for females as well as males (p>0.5).

**Figure 1.**
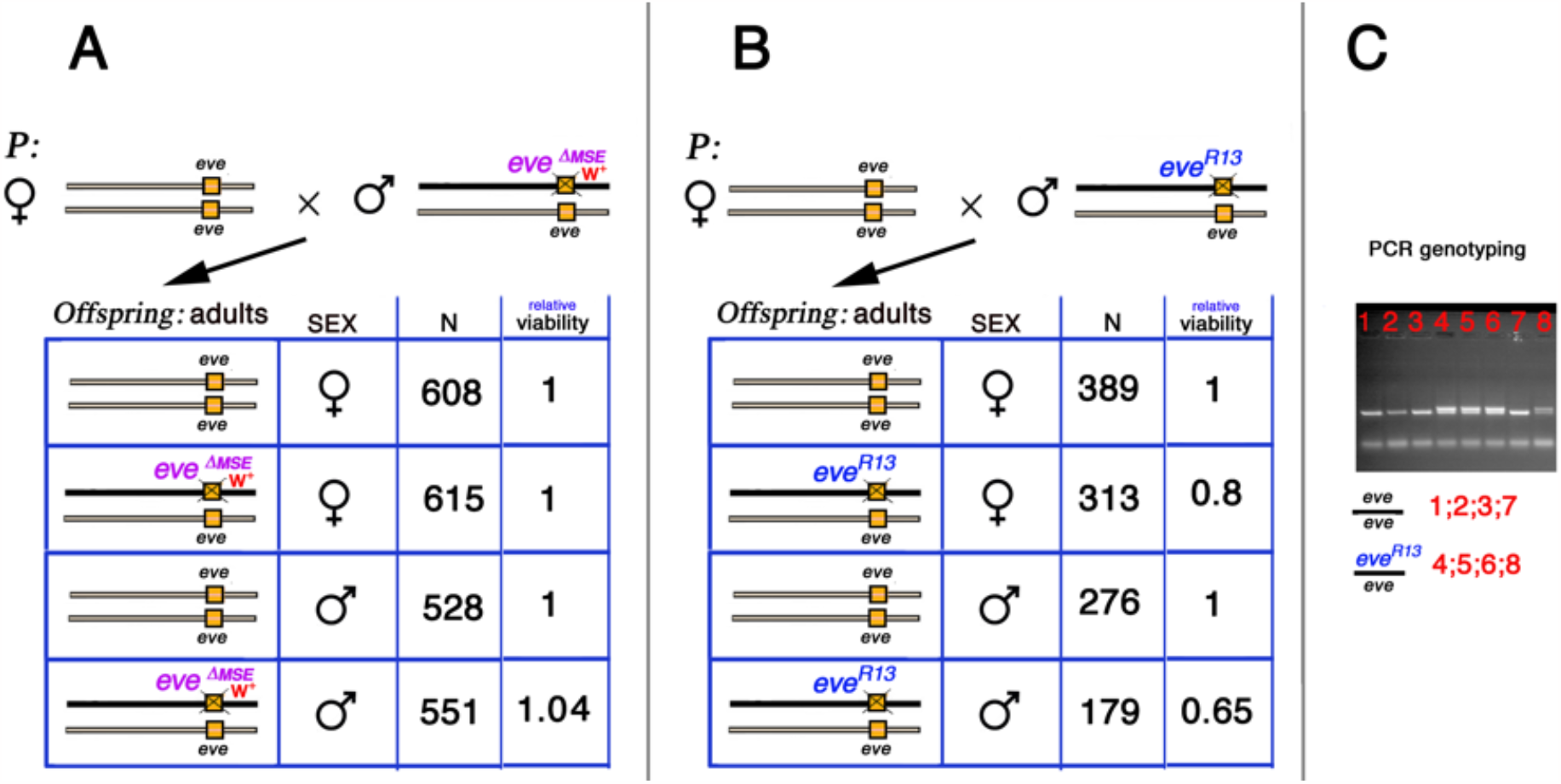
The relative viability of heterozygotes for ***eve*** ^***ΔMSE***^ and ***eve*** ^***R13***^. **A** and **B**. Cross schemas for estimating adult relative viability in ***eve*** ^***ΔMSE***^ and ***eve***^***R13***^ experiments. **C**. Offspring adults were scored by PCR genotyping in ***eve***^***R13***^ experiment.

The relative viability of female and male heterozygotes for ***eve*** ^***R13***^ were 0.8 and 0.65 correspondingly (Figure 1B). The values were significantly different for both females 0.8 (p= 0.0041) as well as males 0.65 (p=0.0001) in comparison to those of females and males not carrying a lethal (viability of these = 1).

### Engrailed (En) patterning in embryos of *eve* ^*R13*^ and *eve* ^*ΔMSE*^ heterozygotes

By demonstrating viability loss only in ***eve*** ^***R13***^ heterozygote we proceeded to investigate whether specific defects in segmentation could be observed in the ***eve*** ^***R13***^ and ***eve*** ^***ΔMSE***^ heterozygote embryos (Figure 2). The EVE stripe 2 developmentally corresponds to parasegment 3, which is bordered by En stripes 3 and 4. It is why we focused attention on the position of En stripe 4 by measuring its location relative to bordering stripes 3 and 5 (Figure 2C.) in the embryos at stage 10 and 11. The ***eve*** ^***R13***^ and ***eve*** ^***ΔMSE***^ mutant embryos were identified by PCR genotyping after image taken. Ratios of parasegment 3 length /3+4 length were 0.5, 0.47, and 0.45 for wildtype, ***eve*** ^***ΔMSE***^, and ***eve***^***R13***^ correspondingly. Both heterozygote genotypes exhibited a statistically significant reduction of parasegment 3 according to the Mann-Whitney Wilcoxon test (Figure 2D and 2G; *p<0*.*001*). In addition, we observed a statistically significant difference of parasegment 3 in the ***eve*** ^***R13***^ and ***eve*** ^***ΔMSE***^ heterozygote embryos according to the Mann-Whitney Wilcoxon test (*p<0*.*01*).

**Figure 2.**
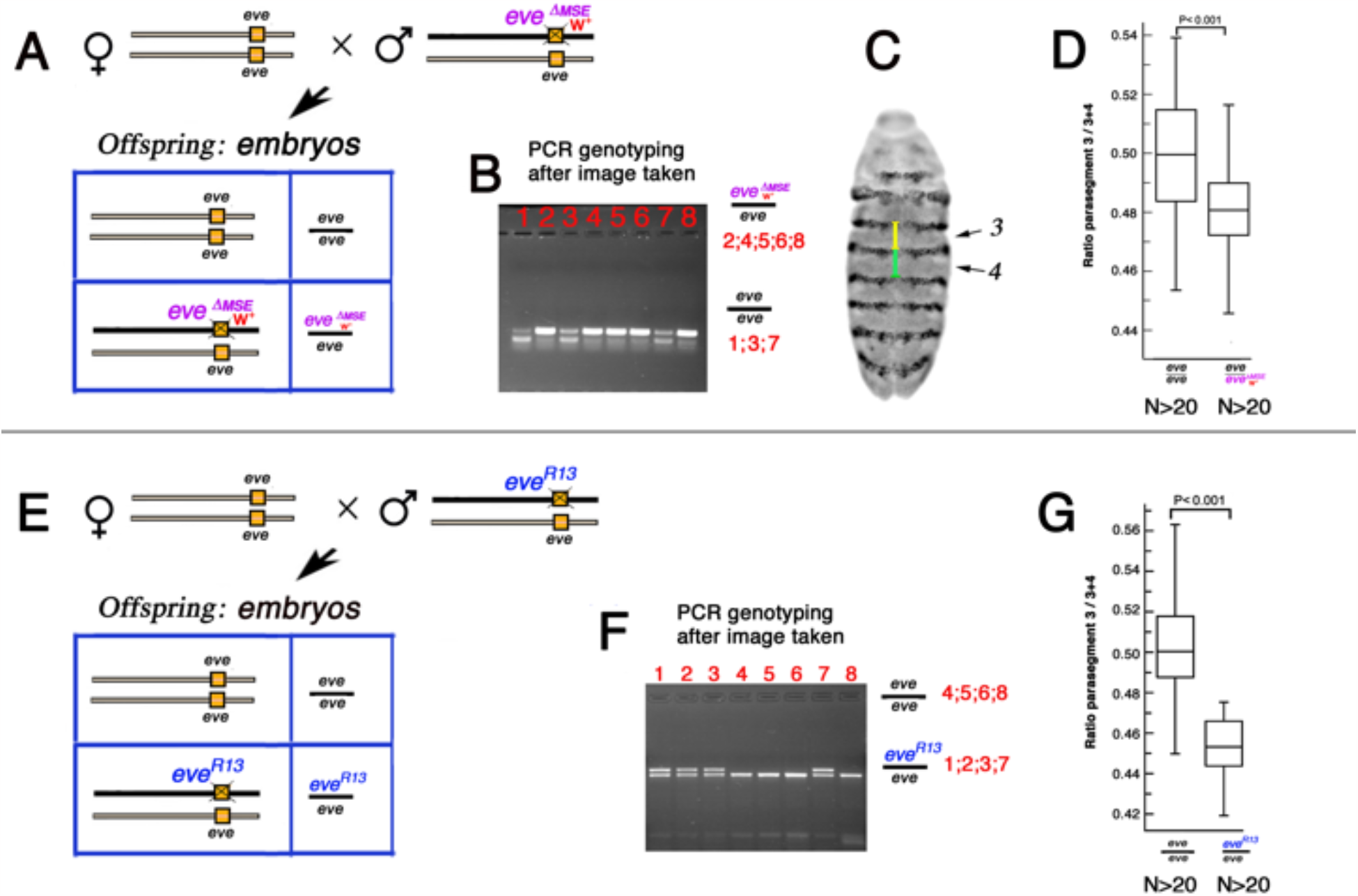
**Engrailed** patterning in ***eve*** ^***R13***^ *and* ***eve*** ^***ΔMSE***^ heterozygotes. Parasegment 3 is reduced in embryos of both ***eve*** ^***R13***^ *and* ***eve*** ^***ΔMSE***^ heterozygotes.**A** and **E**. Cross schema; **B** and **F**. The ***eve*** ^***R13***^ and ***eve*** ^***ΔMSE***^ individual embryos were identified by PCR genotyping after the **En** image taken; **C**. The lengths of parasegments 3 and 4 in the **En** pattern were measured at the ventral midline of embryos. Shown by green and yellow bars respectively; **D** and **G**. Parasegment 3 length relative to sum of parasegments 3+4; stages 10+11. The Mann-Whitney Wilcoxon test. **D. *eve*** ^***ΔMSE***^ heterozygotes vs wildtype homozygotes. **G. *eve*** ^***R13***^ heterozygotes vs wildtype homozygotes.

Our studies of segmentation phenotypes of lethal heterozygotes have demonstrated that ***eve*** is dosage-sensitive haploinsufficient gene. What is more, two lethal alleles showed different extent of abnormalities in heterozygotes phenotypes. However, it is not clear how these segmentation disorders affect viability.

## Materials and Methods

To evaluate the relative viability of our lethal heterozygotes we crossed eight healthy females of the genotype ***w***^***1118***^ with 15 healthy males of the genotype ***w***^**1118**^; ***eve***^***R13***^ ***/+*** or ***w***^**1118**^; ***eve***^***ΔMSE***^ */****+*** at 25°C. Parents were transferred to a fresh culture vial every day for 20 days. The emerging adult offspring were collected every day from the culture vials for a period of 10 days for scoring. This approach ensured that mutants with slow development rates were counted. In all experiments with ***eve***^***ΔMSE***^ the F1 adults were scored for the expression “*white*^+^ transgene” in the eyes. In experiment with ***eve***^***R13***^ the F1 adults were scored by PCR genotyping according to sequence results of Fujioka et al, 1999. The ***eve***^***R13***^ DNA lacks Pvu II restriction site in homeodomain. Embryo collection and fixation was as described in Patel,1994. *Drosophila* embryos were histochemically immunostained with anti-Engrailed monoclonal 4D9 at 1:10 dilution was visualized using HRP-DAB enhanced by nickel (Patel, 1994). Primers and PCR genotyping protocol of individual embryos are available on request. Mutations ***eve*** ^***R13***^ and ***eve***^***ΔMSE***^ have been generated in different labs and in different times. Perhaps, these two stocks have different genetic backgrounds. For now, we assume that this fact does not change our experimental conclusions. However, this subject deserves to be carefully investigated in the future.

## Discussion

We herein address a fundamental biological question: How does a lethal mutant at the non-coding DNA such as enhancer site affect fitness when carried over a “normal wild type” allele? The answer would allow to evaluate both the time of existence and the frequency for existence of a lethal mutation in populations from compensatory evolution point of view.

The availability of a homozygous-lethal *cis*-regulatory mutation ***eve***^***ΔMSE***^ in the ***eve*** locus allowed us to estimate the relative viability of the heterozygote where one allele is normal, and the other allele is a homozygous-lethal mutant. To our surprise the viability of ***eve***^***ΔMSE/***^***/* +** heterozygote was equal to the viability of homozygote with two normal alleles. In contrast, it was not the case for the ***eve***^***R13***^ coding region lethal mutant, which significantly reduced the viability of heterozygotes in females and especially in males.

The S2E drives ***eve*** expression in stripe 2 of the “early” ***eve*** seven stripe expression pattern at stage 5, syncytial blastoderm (Small et al, 1992). Then, the autoregulatory *cis-*element drives “late” seven stripe pattern ***eve*** expression (including stripe two) at cellular blastoderm and early gastrulation stages (Jiang et al, 1991). The gene ***eve*** is pleotropic. Perhaps, the ***eve*** dosage alteration affects not only segmentation pathway but also does the whole *Drosophila* development. For instance, as shown in previous work, the one transgenic copy of the whole ***eve*** locus (with an intact copy of the S2E enhancer) could rescue the lethality of ***eve***^***R13***^***/ eve***^***R13***^ mutant to adulthood only partially (Ludwig et al, 2011). However, two copies of such complete ***eve*** locus (with two copies of S2E enhancer) rescue about 80% of the lethality ***eve***^***R13***^***/ eve***^***R13***^ mutant to adulthood in the same transgenic assay (Ludwig et al, 2011).

That assay above could produce some confusions with current results. One copy of functional transgenic ***eve*** locus can rescue only 15-20% lethality of ***eve***^***R13***^/***eve***^***R13***^ mutant but one copy of normal endogenous allele ***eve*** in heterozygote with lethal allele ***eve*** ^***ΔMSE***^ could restore viability equal to that of the normal homozygote. What is the mechanism for such compensation phenomenon? Our explanation relays on the action of transcriptional autoregulation (Jiang et al, 1991; Crews & Pearson, 2009). The EVE TF regulates its own expression level through the autoregulatory *cis*-regulatory module. It could elevate the expression level from one dose to the level sufficient to restore about 100% fitness in the heterozygote with ***eve*** ^***ΔMSE***^ lethal allele.

The scenarios follow. The S2E drives ***eve*** expression in EVE stripe two at syncytial blastoderm stage. To regulate ***eve*** expression through autoregulatory element it takes combinatorial interactions of multiple transcription factors (activators and repressors) including EVE factor itself at cellular blastoderm and early gastrulation stages (Jiang et al, 1991; Fujioka et al, 1996). The EVE is a transcription repressor. A low concentration of EVE repressor within early stripe two could allow additional transcriptional activation of the ***eve*** in the late stripe two through autoregulatory element. This mechanism does not work in the ***eve***^***R13***^ case because one of the two ***eve*** loci produces truncated dysfunctional EVE protein in the heterozygote. In other words, there is not enough EVE protein activity in the stripe 2 to assemble correct transcription complex with other transcription factors, including at the autoregulatory *cis*-element.

Is *D. melanogaster* ***eve*** being a haploinsufficient gene? By widespread definition, <u>haploinsufficiency</u> is the requirement for two wild-type copies of a gene for a normal phenotype in diploids. For haploinsufficient genes, when one copy of a gene is deleted or contains a loss-of-function mutation, the dosage of normal product generated by the single wild-type gene is not sufficient for correct function. Just to follow the definition, the most direct method to detect ***eve*** haploinsufficiency is to investigate phenotype and fitness of the heterozygous deletion or nonsense mutation in one allele. Our studies of Engrailed segmentation phenotypes of lethal heterozygotes have demonstrated support for the idea that ***eve*** is dosage-sensitive haploinsufficient gene (Nüsslein-Volhard et al, 1985; Chen et al, 2019). Two lethal alleles showed different extent of abnormalities in heterozygotes phenotypes. The situation is less clear how these segmentations disorders affect the relative viabilities. The ***eve*** expresses not only in segmentation pathway but also in many other tissues at different time of the development. In case of ***eve*** ^***ΔMSE***^ heterozygote, perhaps, transcriptional networks employ several distinct tactics, including autoregulation feedback, to ensure wildtype outcome. In contrast, ***eve***^***R13***^ shows different impact on relative viability of heterozygote.

The diverse effect of heterozygotes on both viability and phenotype had special attention in evolutionary genetics (Kimura,1983). In one particular experiment, it has been found out that 77 different X-chromosomal lethals have different viability numbers ranging from 0.602 to 1.312 in heterozygotes as compared with lethal-free homozygotes (viability of those=1) (Stern et al, 1952). Most studied heterozygotes were without morphological effect. The lethal mutants had not been molecularly characterized in the time. Half of the mutants had spontaneous origin, while another half of studied mutants was generated by ionizing radiation. Unfortunately, it is not possible to distinguish between coding regions mutants from non-coding region in the (Stern et. al., 1952) experiment.

One of the main results of our study that two deleterious noncomplemented mutations from the same locus have showed different viability in heterozygotes. Whether this discovered phenomenon could be observed in each eukaryotic TF locus? We argue that perhaps, it is. The TF loci employ a multiplicity of mechanisms that contribute to robust gene expression in two allele diploid organism. Expression of one allele with deleterious reduced gene dosage mutation could be compensated with transcriptional network tactics, including autoregulation feedback, by another ‘wild type’ allele, that by itself would not be sufficient to provide enough protein on its own. The functional relationship between phenotypes of the alleles described above is another possible molecular mechanism of genetic dominance.

In conclusion, we show that heterozygotes for the coding region mutant ***eve*** ^***R13***^ and the *cis-*regulatory mutant ***eve*** ^***ΔMSE***^ demonstrate different impact on relative viability and on the embryonic segmentation. A novel hypothetical route for evolution emerges from the presented results. Based on these results we predict that, some *cis-*regulatory mutants (including the deleterious ones) may exist as heterozygotes with high fitness for many generations in populations, before possibly encountering an epistatic interaction with a compensatory mutation at a different site in the genome. This study is consistent with the compensatory mechanisms of evolution for functionally important *cis-*regulatory elements.

## Acknowledgements

Author expresses deep respect and special thanks for collaboration to Dr. Marty Kreitman. The work was supported by the National Science Foundation under award 1916895 to Martin Kreitman, by NIH 2 R01 OD010936-27A1to Dr. John Reinitz, and by NIH 2 RO1 GM127366-03A1 to Urs Schmidt-Ott. I thank Dr. Natalia Tamarina for discussion and comments.

